# Genetic context and proliferation determine contrasting patterns of copy number amplification and loss for 5S and 45S ribosomal DNA arrays in cancer

**DOI:** 10.1101/154120

**Authors:** Meng Wang, Bernardo Lemos

**Author notes:** Correspondence to: Bernardo Lemos Department of Environmental Health & Molecular and Integrative Physiological Sciences program Harvard T. H. Chan School of Public Health 665 Huntington Avenue, Boston, MA 02115, Bldg 2-219, Boston, MA, USA.

## Abstract

The multicopy 45S ribosomal DNA (45S rDNA) array gives origin to the nucleolus, the first discovered nuclear organelle, site of Poll I 45S rRNA transcription and key regulator of cellular metabolism, DNA repair response, genome stability, and global epigenetic states. The multicopy 5S ribosomal DNA array (5S rDNA) is located on a separate chromosome, encodes the 5S rRNA transcribed by Pol III, and exhibits concerted copy number variation (cCNV) with the 45S rDNA array in human blood. Here we combined genomic data from >700 tumors and normal tissues to provide a portrait of ribosomal DNA variation in human tissues and cancers of diverse mutational signatures. We show that most cancers undergo coupled 5S rDNA array amplification and 45S rDNA loss, with abundant inter-individual variation in rDNA copy number of both arrays, and concerted modulation of 5S-45S copy number in some but not all tissues. Analysis of genetic context revealed associations between the presence of specific somatic alterations, such as P53 mutations in stomach and lung adenocarcinomas, and coupled 5S gain / 45S loss. Finally, we show that increased proliferation rates along cancer lineages can partially explain contrasting copy number changes in the 5S and 45S rDNA arrays. We suggest that 5S rDNA amplification facilitates increased ribosomal synthesis in cancer, whereas 45S rDNA loss emerges as a byproduct of transcription-replication conflict in highly proliferating tumor cells. Our results highlight the tissue-specificity of concerted copy number variation and uncover contrasting changes in 5S and 45S rDNA copy number along rapidly proliferating cell lineages.

**Lay Summary:** The 45S and 5S ribosomal DNA (rDNA) arrays contain hundreds of rDNA copies, with substantial variability across individuals and species. Although physically unlinked, both arrays exhibit concerted copy number variation. However, whether concerted copy number is universally observed across all tissues is unknown. It also remains unknown if rDNA copy number may vary in tissues and cancer lineages. Here we showed that most cancers undergo coupled 5S rDNA array amplification and 45S rDNA loss, and concerted 5S-45S copy number variation in some but not all tissues. The coupled 5S amplification and 45S loss is associated with the presence of certain somatic genetic variations, as well as increased cancerous cell proliferation rate. Our research highlights the tissue-specificity of concerted copy number variation and uncover contrasting changes in rDNA copy number along rapidly proliferating cell lineages. Our observations raise the prospects of using 5S and 45S ribosomal DNA states as indicators of cancer status and targets in new strategies for cancer therapy.

## Introduction

The ribosomal DNA (rDNA) arrays give origin to the nucleolus, the nuclear organelle that is the site of ribosomal RNA (rRNA) transcription and ribosome biogenesis [1]. The rRNAs constitute the vast majority of cellular RNAs and are encoded from two kinds of tandemly repeated ribosomal DNA (rDNA) arrays [2-5]. The 45S rDNA array is localized on five human chromosomes, is transcribed by Pol I, encodes the 45S rRNAs that are processed into three rRNAs (18S, 5.8S and 28S rRNAs), and organizes the formation of the nucleolus [6]. The 5S rDNA array is exclusive to human chromosome 1, encodes the Pol III transcribed 5S rRNA, and localizes, together with dispersed tRNAs, at the periphery of the nucleolus. Stunningly high transcription of the rRNAs is required to supply ribosomes, essential cellular machines with tightly controlled and strikingly complex biogenesis involving products transcribed by all three RNA polymerases (Pol). Indeed, the human ribosome is composed of about 80 cytoplasmic ribosomal proteins (cRP) and four ribosomal RNAs (rRNA; 18S, 5.8S, 28S, and 5S components), responsible for protein production and the translation of all protein-coding mRNAs. Modulation of rRNA synthesis during the cell cycle is critical for cell growth and proliferation [7-9]. To initiate rRNA transcription, proteins such as UBF and TIF are required to facilitate Pol I binding onto the rDNA promoter – a region that includes the upstream control element (UCE) and the core sequence. The protein components of the cytoplasmic ribosomes, cRPs, are co-expressed to ensure stoichiometric balance [10-15]. Moreover, hundreds of other proteins and several snoRNAs are needed for the cleavage, modification, transport, and assembly of rRNAs, and the maturation of ribosome in the nucleolus and nucleoplasm.

It is thus expected that altered ribosomal biogenesis and rRNA regulation had been linked to human diseases. The dysregulation of cRP genes (cRPGs) can destabilize rRNAs and/or disturb their synthesis [16, 17], or alternatively participate in tumorigenesis through P53-related pathways [18, 19]. Notably, many well established oncogenes and tumor-suppressor genes are directly or indirectly involved in regulating ribosomal biogenesis and/or nucleolus function [20-22]. For example, proteins in the retinoblastoma (RB) family as well as P53 and MDM2/4 regulate nucleolar function and rRNA synthesis: they can restrict rRNA synthesis by interacting with UBF and repressing Pol I activity [23-25]. Also, the proto-oncogene, c-Myc, can target cRPGs as well as other nucleolar proteins such as nucleophosmin (NPM) and nucleolin [26-28]. Collectively, the observations imply an integrated network of cellular functions that is coherently modulated and is centered on rDNA/nucleolus maintenance, rRNA expression, and ribosome biogenesis.

Both the 5S and 45S rDNA arrays display remarkably variable copy number (CN), ranging from tens to hundreds of copies among eukaryotes [3, 5, 29-32] and displaying a 10-fold variation among individuals in human populations [3, 5, 30]. Surprisingly, copy number of the 5S and 45S rDNA arrays in human blood is positively correlated across genotypes in spite of 5S and 45S array location in separate chromosomes and lack of sequence homology [3]. The observations indicated that copy number in both arrays is jointly modulated and suggested a general mechanism of concerted copy number variation (cCNV) that is likely to operate in other tissue types to maintain balanced 5S and 45S rRNA. Notwithstanding, only a fraction of the rDNA units are transcribed per cell [33, 34] and alternative mechanisms exist to modulate 45S rRNA supply. In addition, rDNA CN itself might contribute to nucleolar function, providing a critical mechanism to maintain genome stability [35] and exerting genome-wide consequences to gene regulation [30, 36]. Finally, a number of seminal ultra-structural studies documented alterations in nucleolar morphology during carcinogenesis [37-40]. In spite of the manifold cellular roles of the rDNA array, the manifestation of concerted copy number variation of the 5S and 45S arrays has not been examined in human tissues other than blood. Furthermore, the role of rDNA CN and altered ribosomal function in human cancers remain crudely characterized. Here we ascertained rDNA copy number in 6 tissues from hundreds of individuals and cancer genomes. We apply corrections for aneuploidy and sequencing batch to provide a comprehensive portrait of rDNA variation in cancer and normal tissues. The data is integrated with genetic context to reveal associations between specific somatic alterations and somatic rDNA CN changes, as well as the impact of proliferation rate.

## Results

### Estimating rDNA copy number in human cancers

Ribosomal RNAs are transcribed from the 45S and 5S rDNA repeats, both of which display ~10-fold variation in copy number within populations. Here we used whole-genome sequencing data (WGS) from the TCGA to ascertain rDNA CN variation across 946 samples (721 tumor and 225 adjacent normal tissue) representing six cancer types with the largest numbers of individuals with paired tumor and adjacent normal tissue [i.e., bladder urothelial carcinoma (BLCA), lung adenocarcinoma (LUAD), lung squamous cell carcinoma (LUSC), kidney renal clear cell carcinoma (KICA), head and neck squamous cell carcinoma (HNSC), and stomach adenocarcinoma (STAD)] (Table 1). We adopted a computational approach developed in earlier studies [3, 30] with important modifications to estimate rDNA CN in tumors. Briefly, rDNA CN was estimated by dividing the average depth of each component (18S, 5.8S, 28S and 5S) by the average depth of the selected background, i.e. single copy exons and introns residing on chromosomes bearing rDNA arrays. To diminish the effect of heterogeneity within 18S and 28S sequences, we identified segments with the smallest coefficient of variation in read depth across sites to represent the two components (Figure 1A). Furthermore, tumor aneuploidy is rampant [41, 42] and can be a major confounder in rDNA CN estimates. Hence, we accounted for ploidy variation in tumors using estimates of per gene copy number amplification/loss from the FireBrowse portal, which were ascertained by the GISTIC 2.0 pipeline [43] using the genome-wide SNP array data. We confirmed extensive ploidy changes in tumors (Figure S1), which suggested the need for correction in rDNA CN estimates. Tumor aneuploidy and gene amplification/loss will reflect on the segment sequencing depth, such that a positive correlation between ploidy estimates and read depth is expected. Indeed, we did observe such relationship in tumor samples (Figure S2A), which supported our methodology and underscored the necessity of correction for aneuploidy. Intriguingly, we also detected significant batch effects of TCGA plate ID on estimates of rDNA CN (Figure S2B, Table S1), which are reminiscent of batch effects observed in recent mtDNA copy number estimates for tumors [44]. Here we specifically controlled for sequencing batch using linear regression model separately for each cancer type (see Methods). Finally, we identified rDNA reads by “slicing” of (BAM) files generated by the TCGA consortium (see Methods). The approach had a marginal effect on rDNA estimates, when compared to *de novo* identification of rDNA reads by processing and mapping of raw reads to rDNA references (Figure S2C).

**Table 1.** Cancer types and sample sizes with whole genome DNA sequencing data (N = 721 patients).

| Cancer type | TCGA ID | Number of tumor samples |
| --- | --- | --- |
| Bladder Urothelial Carcinoma | BLCA | 113 |
| Lung adenocarcinoma | LUAD | 233 |
| Lung squamous cell carcinoma | LUSC | 50 |
| Stomach adenocarcinoma | STAD | 127 |
| Kidney renal clear cell carcinoma | KIRC | 44 |
| Head and Neck squamous cell carcinoma | HNSC | 154 |

**Figure 1.**
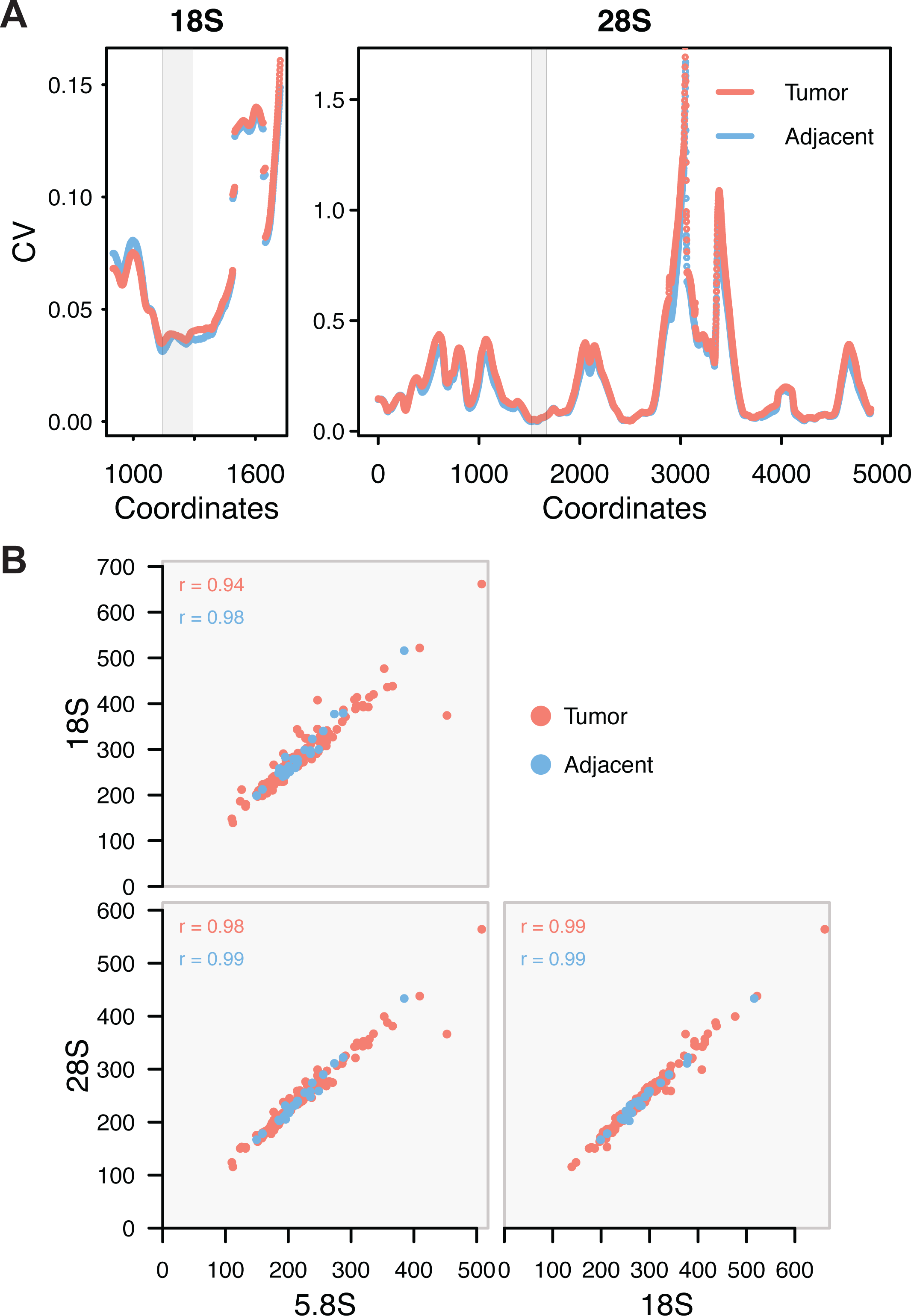
Estimating rDNA copy number. (A) For each 150bps window along the rDNA array reference, we calculated the coefficient of variation (CV) of the average depth among samples. The x-axes indicate the starting coordinate for each window on 18S (901-1871) and 28S (whole length). LUAD tumor and adjacent tissues have largely consistent CV across 18S and 28S sequences. The grey regions highlight the windows selected to assess CN in the 18S and 28S components. (B) Nearly perfect pairwise correlations between 45S components. BLCA is displayed as an example. Copy number estimates are corrected for batch and ploidy.

Due to their close physical linkage, dosage (copy number) of the 45S rDNA components (18S, 5.8S and 28S rDNA) is expected to be congruent with each other, although not necessarily identical. Indeed, we observed positive correlations among 18S, 5.8S and 28S components in normal adjacent tissue of patients with all cancer types (Figure 1D and S3; Pearson’s r = 0.47 − 0.99, P < 0.005 for all samples). However, rDNA components displayed lower correlations in tumors, likely due to ploidy variation of background genes and/or higher prevalence of non-canonical and truncated rDNA units [45, 46]. Accordingly, upon removal of aneuploidy and sequencing batch confounders, we observed that the 45S components had near perfect correlations among each other in LUAD, STAD, BLCA, and HNSC (Figure 1B and S3; P < 0.0001 for all samples). The analyses improved on earlier methods for computing rDNA CN estimates [30] and show that rDNA CN can be reliably ascertained in both cancer and normal tissues.

### Concerted copy number variation is not universally manifested in all tissues

The 5S array is located on an unrelated chromosome and has evolved copy number variation that is tightly correlated with copy number of the 45S rDNA in B-cell derived lymphoblastoid cell lines (LCLs) and whole blood [3]. Here we applied the improved methodology and corrections to ascertain rDNA copy number of the 5S rDNA and address whether the 5S and 45S rDNA arrays display cCNV in human solid tissues. We observed that the manifestation of cCNV is variable among rDNA components and across tissues (Figure 2 and S3). Specifically, we observed strong cCNV for both tumor and adjacent tissues in BLCA and LUAD, although only the 5.8S is significantly correlated with 5S in KIRC and LUSC. On the other hand, cCNV appeared absent in HNSC and STAD. These results reveal that the strength of cCNV is variable across tissues, and suggest that the phenomenon is not universally manifested in all tissues.

**Figure 2.**
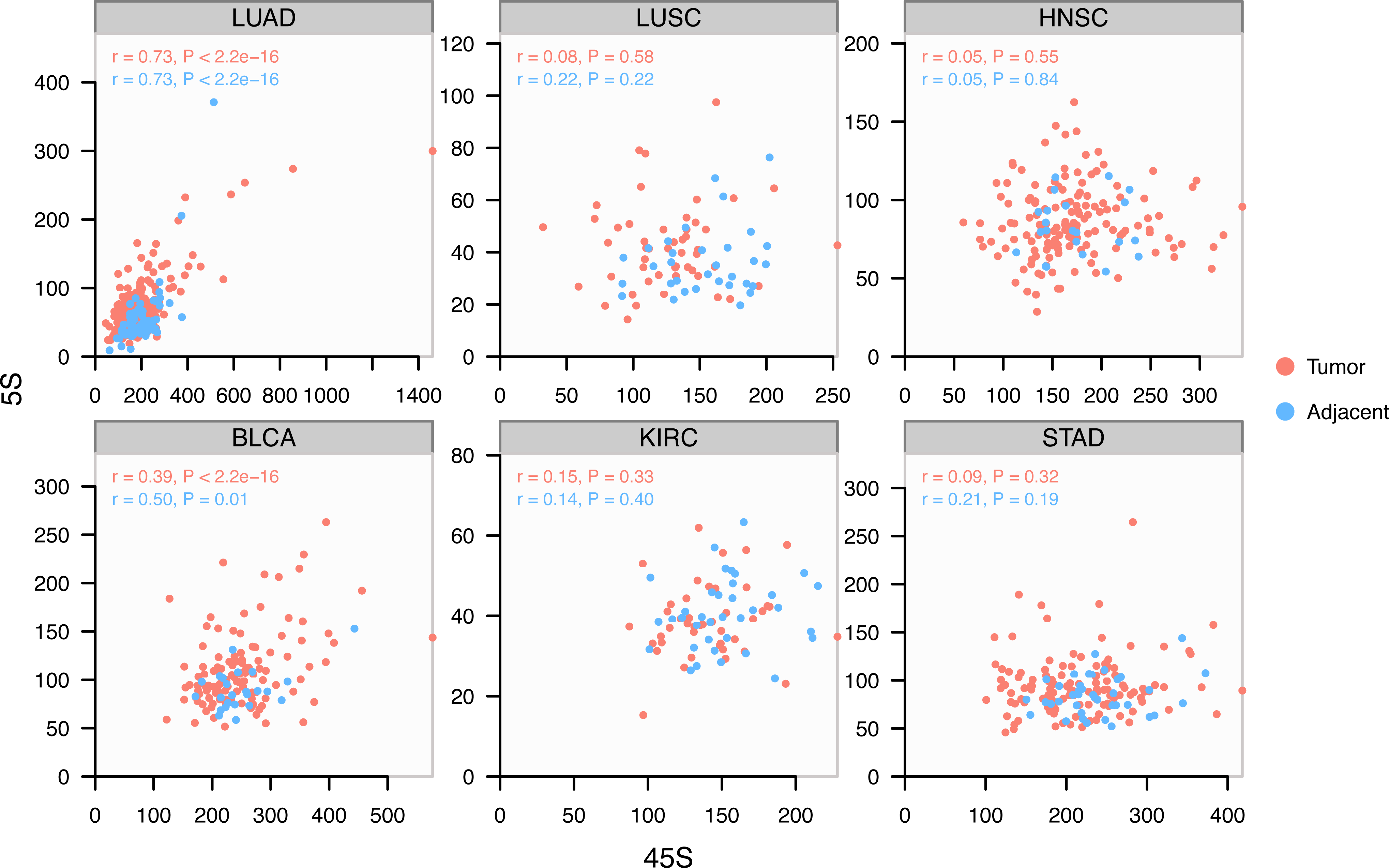
Variable manifestation of concerted copy number variation (cCNV) across tissues. Correlation between the 5S and 45S rDNA arrays is variable among tissues. The correlation is significant and similarly manifested in normal and cancer lineages of LUAD and BCLA.

### Ribosomal DNA amplification and loss in paired cancer and adjacent normal tissue

We compared paired tumor and adjacent tissue within an individual to obtain estimates of rDNA fold-change amplification and loss in cancer lineages. To minimize confounders due to plate ID, analyses with tumor-adjacent comparisons only included 225 patients with both tumor and adjacent tissue sequenced in the same plate. Intriguingly, we found a mild but significant depletion of 45S rDNA in 5 out of 6 cancer types relative to adjacent normal tissue (Figure 3). The exception is BLCA showing only a slight but not significant reduction in 45S rDNA. The events of 45S loss are especially salient in LUAD with 73% (54/74) of all paired-adjacent contrasts revealing loss events, with 13% (7/54) of all loss events less than 60% of the copy number values in the adjacent tissue. On the other hand, we observed, amplification of the 5S rDNA array in 4 cancer types, regardless of loss in the 45S components (Figure 3). Interestingly, LUAD stood out with 68.9% (51/74) of the cases showing amplification events, with 47.1% (24/51) of them reflecting a >40% increase in 5S rDNA copy number. Notably, cCNV between 5S and 45S in LUAD tumors remains detectable, in spite of their contrasting patterns of alterations. We conclude that 5S rDNA amplification is a recurrent event in cancer lineages and is frequently accompanied by loss of 45S array components.

**Figure 3.**
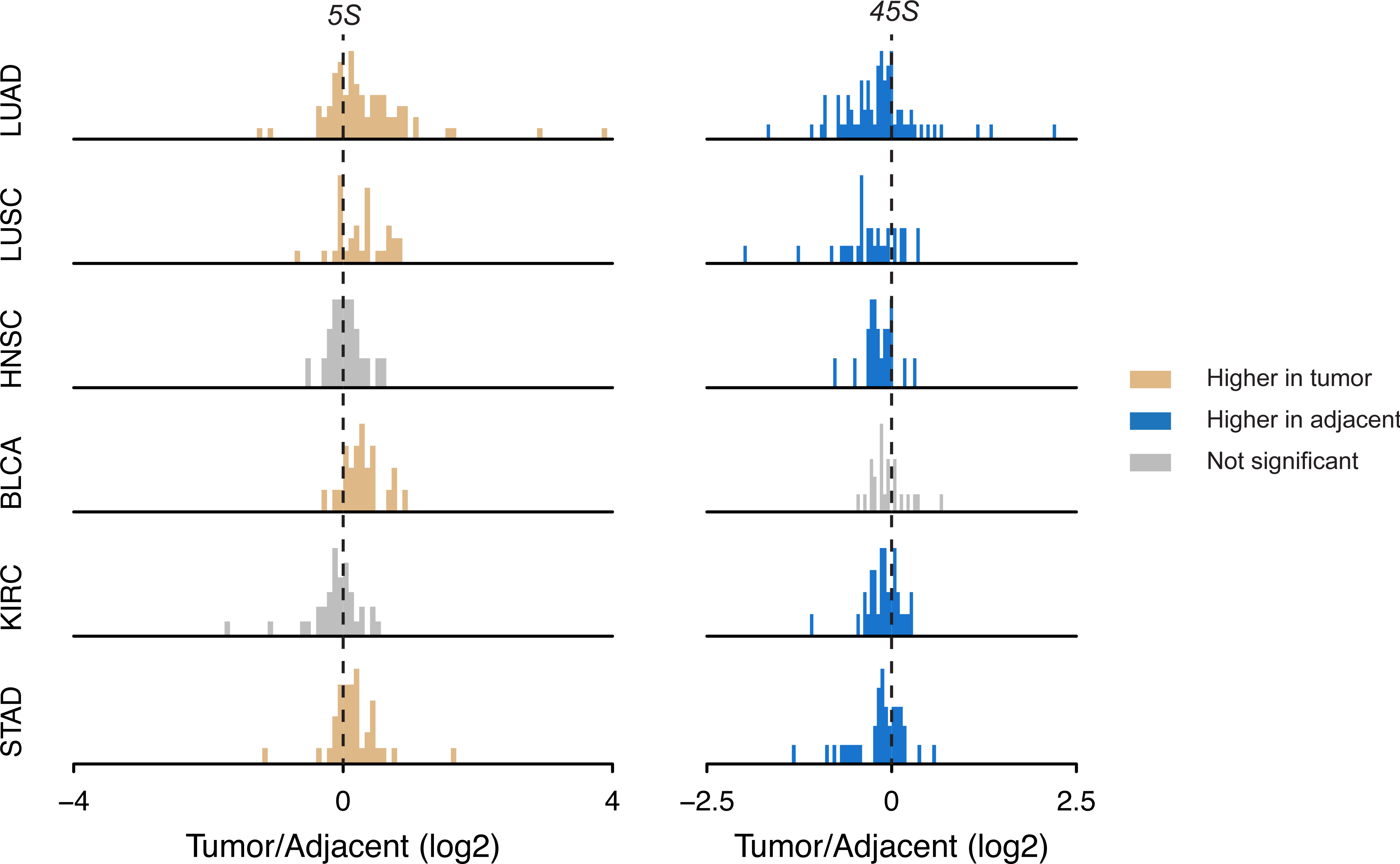
rDNA CN amplification and loss between cancer tissue and paired adjacent normal tissue. Yellow and blue denote significant gain or loss in tumors compared with paired adjacent controls (Wilcoxon rank sum test P < 0.01). There are 74, 32, 22, 24, 35 and 38 informative patients for LUAD, LUSC, HNSC, BLCA, KIRC and STAD respectively.

### Cancer somatic copy number alterations (SCNAs) couple 5S expansion and 45S loss

Multiple mutations are involved in cancer development. Notably, mutations in genes that control genome stability can trigger numerous downstream mutation events [42, 47, 48]. Therefore, we investigated the impact of genomic variations on rDNA CN in tumors. To address the issue, we cross-referenced our estimates of rDNA CN with estimates of local SCNAs as ascertained by the TCGA consortium. The data allowed us to stratify tumors by presence or absence of a SCNA, and examine their association with CN of 5S or 45S. As a result, we observed 39 and 38 events significantly correlated with 5S and 45S, respectively, in at least one cancer type (Figure 4A, P < 0.05, Table S2). Reassuringly, the strongest association for 5S is its positive correlation with 1q42.3 amplification in STAD (Figure 4A). This association is expected because the 5S rDNA resides within the 1q42 segment. Besides 1q42 amplification, most other significant rDNA-associated SCNAs are physically unlinked to either the 45S or the 5S rDNA array. For example, 9q34.3 amplification is significantly associated with the accumulation of 5S (Figure 4A, linear regression’s coefficient = 31.15 and P = 5.23e-6), whereas 15q11.2 deletion is associated with 45S loss in STAD (Figure 4A, coefficient = -91.86 and P = 0.00022). Most notably, the majority of the significant SCNA-45S associations are implicated in greater 45S loss along the cancer lineages (Figure 4B). Specifically, 9/14 SCNAs amplification events and 23/25 SCNAs deletion events resulted in greater 45S rDNA loss (binomial test for total, P = 7.025e-05). On the other hand, the majority of significant SCNA-5S associations are implicated with events of 5S amplification along the cancer lineage (Figure 4B). Specifically, 15/19 SCN amplification events and 11/19 SCN deletion events resulted in greater 5S amplification (P = 0.034 for total). Because of the frequent cooccurrence of 5S gain and 45S loss, we correlated SCNAs with 5S / 45S ratios. In agreement with our expectations, we found that nearly all of the statistically significant rDNA-SCNA associations are implicated in higher 5S/45S ratios in the tumor lineage (Figure 4B, 71/81, P = 1.799e-12), regardless of whether the focal SCNA was a deletion or a duplication event.

**Figure 4.**
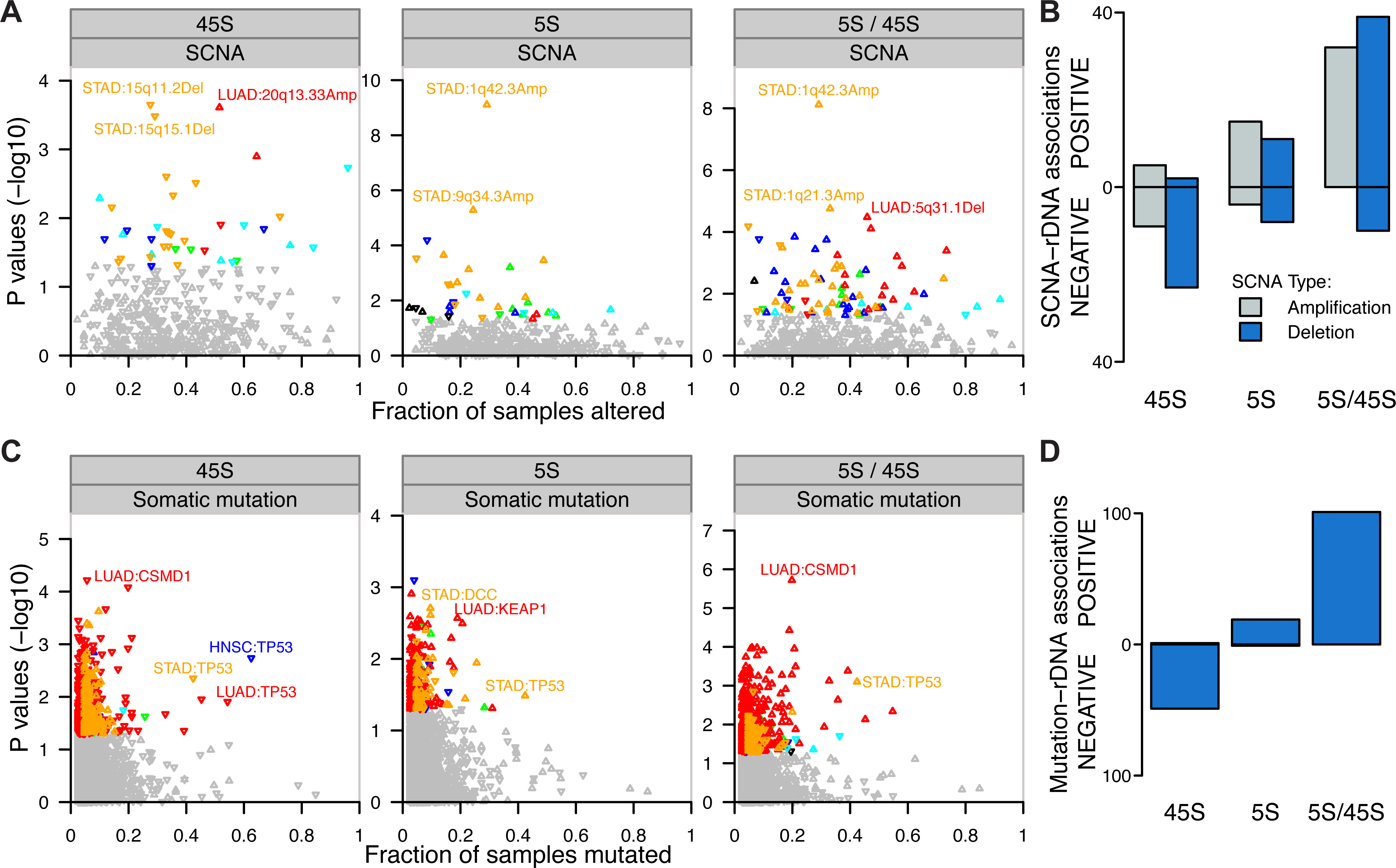
Association between genetic context and rDNA CN alterations. (A) Associations between copy number alterations (SCNA) and 5S CN, 45S CN and 5S/45S ratio. (B) Significantly associated rDNA-SCNA pairs (P < 0.05) are preferentially implicated in greater 45S loss and greater 5S rDNA amplification (P < 0.05, binomial test). (C) Association between somatic mutations and 5S CN, 45S CN and 5S/45S ratio. (D) Significantly associated mutation-rDNA pairs (P < 0.05) are almost exclusively implicated in greater 45S loss and greater 5S rDNA amplification (P < 0.001, binomial test) in LUAD. For (B, C) Y-axis show the P-values for the associations between the SCNA or gene mutation event and 45S CN, 5S CN and 5S/45S ratio. rDNA associations were colored according to cancer type (P < 0.05). The up/down direction of triangles indicates that somatic alteration is associated with increased or decreased CN or 5S/45S ratio. The X-axis shows the fraction of patients with the focal SCNAs (ploidy cutoff > 2.1 for amplification and < 1.9 for deletion) or non-silent gene mutation.

## Dosage of the 5S rDNA is increased through 1q42 segmental duplication but also array expansion

To further understand the mechanisms for 5S amplification, we used two genes localized immediately upstream and downstream of the 5S array (RNF187 and RHOU), as a proxy to address segmental duplications involving the 1q42.13 region. First, we observed significant positive correlations between estimates of 1q42.13 ploidy and 5S rDNA CN in LUAD, LUSC and STAD (Figure 5). Second, we observed that the 1q42.13 segment is significantly amplified in all but one cancer type (Figure 6A). This suggests that amplification of the 1q42 segment is a common and recurrent mechanism causing increased 5S CN across distinct cancers. On the other hand, recombination within the 5S rDNA array could provide an alternative mechanism contributing to 5S rDNA amplification. To address the issue we examined patients for which ploidy estimates at 1q42.13 is closest to diploidy (1.98-2.02). Interestingly, significant 5S amplification was still observed in these individuals for which 1q42 ploidy did not increase (Figure 6B, one-tailed Wilcoxon rank sum test, P = 0.0014, STAD). In conclusion, the data suggest that both 1q.42 segmental duplications as well as local 5S rDNA array expansion contribute towards increased 5S rDNA dosage in cancer lineages.

**Figure 5.**
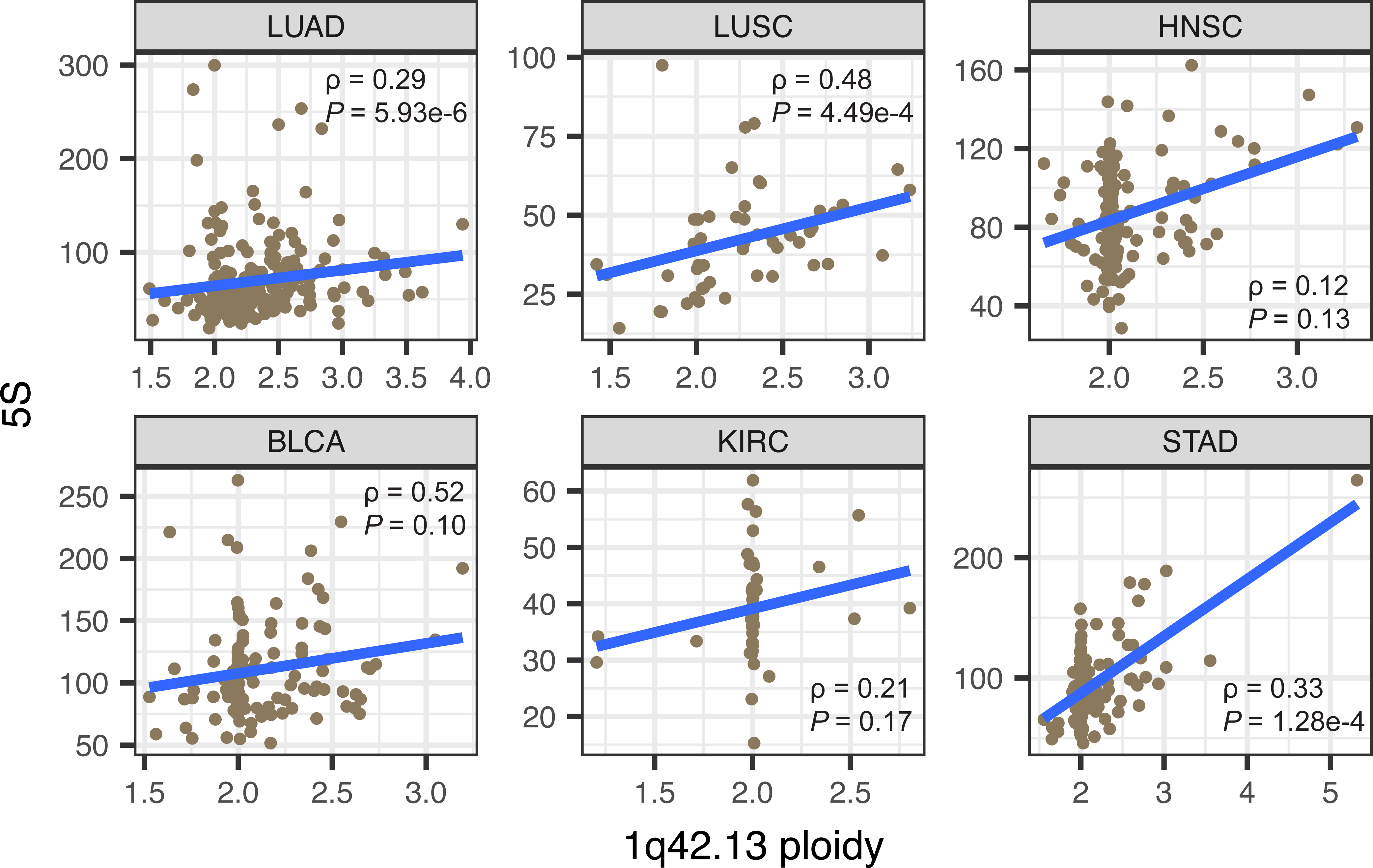
Increased 1q42.13 ploidy partially explains increased 5S rDNA CN in cancers. In each cancer type, all available tumors were included. Spearman’s rank correlation was used.

**Figure 6.**
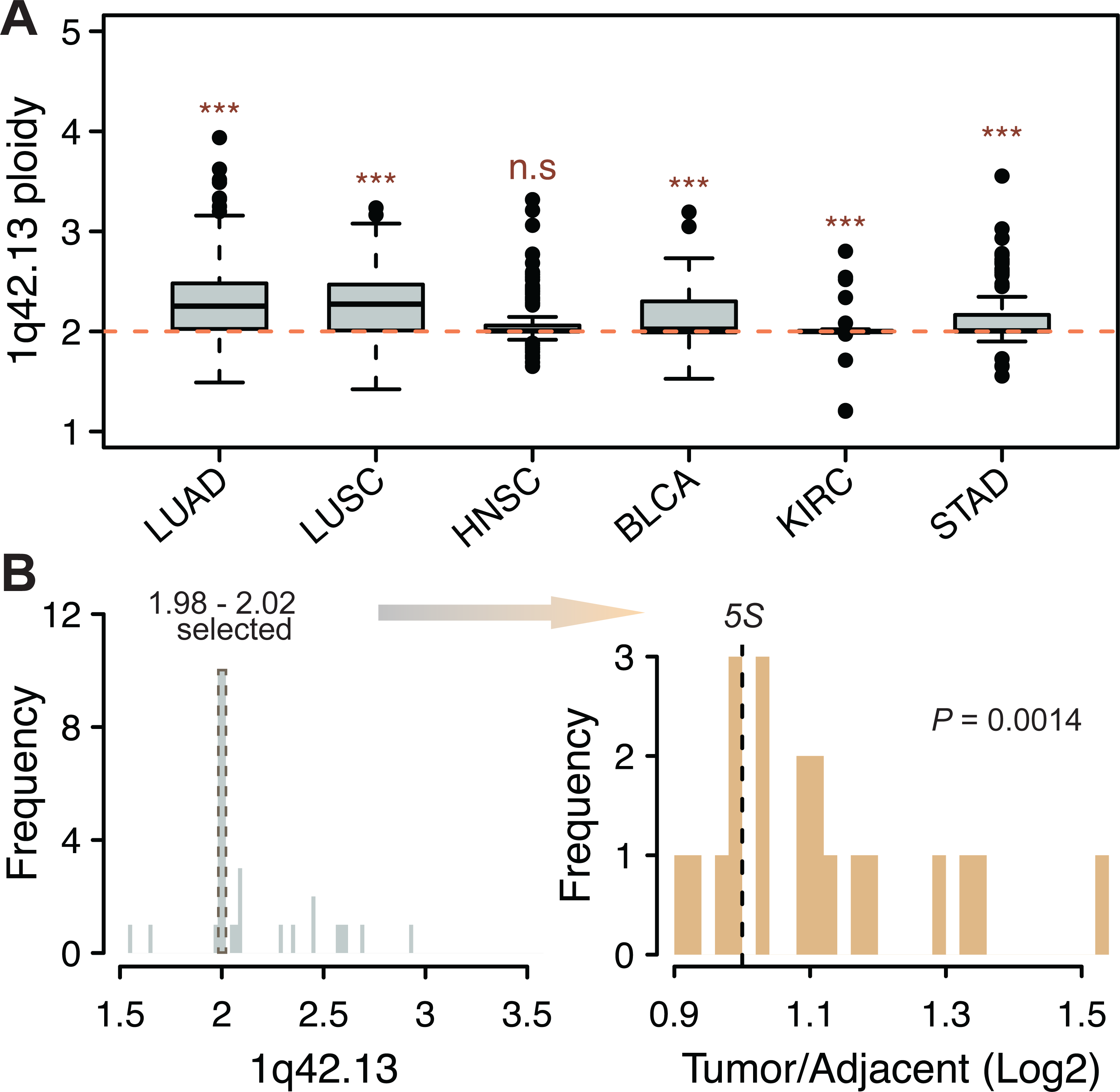
The 5S rDNA is increased through 1q42 segmental duplications and 5S array expansion. (A) Most cancers displayed significantly increased 1q42.13 ploidy. (B) Significant 5S CN amplification was still observed in STAD when only considering patients that are closest to diploidy at 1q42.13 (P-value from one-tailed Wilcoxon rank sum test). Only patients with 5S rDNA CN estimated for both tumor and adjacent tissues are shown in B.

### Cancer P53 mutations couple 5S expansion and 45S loss

We next examined somatic mutations associated with rDNA CN amplification or loss in each cancer type. Specifically, we identified tumors containing non-silent somatic mutation(s) in protein-coding genes and compared their rDNA CN amplification or loss events with those tumors from individuals without cancer mutations in the focal gene. We examined a total of 17,035 genes containing somatic mutations with 1,481 of which being present in ten to a few hundred individuals in at least one cancer type. Strikingly, we found that, TP53 mutations are negatively associated with 45S CN in STAD, HNSC (P < 0.005, Wilcoxon rank sum test) and LUAD (P = 0.012, Wilcoxon rank sum test) (Figure 4C and Table S3). On the other hand, most LUSC tumors show TP53 mutations while most KIRC tumors do not show TP53 mutations, leading to low statistical power for TP53-rDNA association in both cases. Presence of TP53 mutation is also associated with 5S amplification in STAD (Figure 4C, P = 0.033). Overall, the associations between mutation presence and rDNA CN are particularly apparent in LUAD, the cancer with the largest sample size. In this case, the associations between mutation presence and 45S rDNA CN are predominantly negative (Figure 4D, 49/50, binomial test P = 9.059e-14), whereas the association between mutation presence and 5S CN are predominantly positive (19/20, P = 4.005e-5). The positive associations for the 5S rDNA reinforce the notion that tumors might be selected for 5S rDNA CN increases, possibly to supply the cell with increased protein synthesis and proliferative capacity. Finally, the contrasting direction of the gene-associations with 5S and 45S are most evident when examining the ratio of 5S / 45S rDNA in the presence of somatic mutations: all 101significant gene-rDNA (5S/45S ratio) associations are positive (P < 2.2e-16). This indicates a higher 5S/45S ratio in the presence of the mutation. To increase statistical power, we combined all tumors in a pan-cancer analysis of gene-rDNA association. For each gene, we applied an ANOVA to compare 45S, 5S or their ratio between the groups with and without the mutation, with cancer types as a covariate. We observed dozens of genes significantly associated with 45S, 5S, and their ratio, respectively (Table S4). In this analysis TP53 mutations emerged again as one of the top candidates associated with significantly lower 45S (P = 0.0015), as well as significantly higher 5S / 45S ratio (P = 5.24e-9). Among genes associated with the 5S we find that 89.5% (128/143) of them displayed positive associations, indicating higher 5S rDNA CN in the presence of the mutation. Finally, in this pan-cancer analysis, >95% of all significant associations between the presence of the somatic mutation and the 5S / 45S ratio are positive.

### Increased proliferation rates as drives of coupled 5S rDNA amplification / 45S rDNA loss

Tumors undergo increased ribosome biogenesis and accelerated proliferation rates [49]. We hypothesize that contrasting rDNA CN variation may reflect underlying rapid tumor cell proliferation. One hypothesis is that increased proliferation is facilitated by greater 5S rDNA dosage. A related hypothesis is that 45S rDNA loss emerges as a byproduct of transcription-replication conflict in rapidly proliferating cells. To address the issues we calculated a proliferation index (PRI) across tumors and adjacent tissues. PRI is based on the expression of 793 genes (denoted as ‘YW’ set) which were recently identified by positive correlation with proliferation rate across 60 cancer cell lines [50]. A larger PRI indicates higher proliferation in the cell lineage. We also measured PRI using a second set of 350 genes implicated in cell proliferation rates [51] (denoted as ‘RS’ set). While the YW and RS sets are mostly independent of each other (only 74 genes are shared), we observed highly positive correlations between PRI estimates using the two sets (Figure S4, Spearman’s Rho = 0.53-0.89, P < 2.2e-16). We compared tumor tissues relative to their adjacent normal, and indeed observed significantly increasing PRI for almost all cancer types (Figure S5) with LUSC, LUAD, and STAD among the top cancers showing the largest PRI increases in cancer relative to adjacent normal. Notably, higher PRI is also associated with lower survival probability (Figure S6A and S6B, also in [50]), as well as more advanced tumor stage (Figure S6C and S6D).

Next, we focused on LUAD, the cancer with the highest number of adjacent-tumor pairs, to examine whether tumor proliferation can explain changes in rDNA CN. In agreement with our hypothesis, we observed that 45S rDNA CN is negatively correlated with PRI (Fig 7A, rho = -0.14, P = 0.037; Fig S7A), whereas 5S rDNA CN is positively correlated with PRI (Fig 7A, rho = 0.12, P = 0.076; Fig S7A). The 5S / 45S ratio is also positively correlated with PRI (Fig 7B, Spearman’s rho = 0.23, P < 0.001; Fig S7A). To verify the pattern, we then selected individuals with rDNA CN and expression data available in both tumor and adjacent tissues. For these individuals, we calculated the relative fold change (FC) in PRI in tumor relative to its adjacent tissue, and correlated it with the FC in 5S rDNA CN, 45S rDNA CN or their ratio for the same patients. Thirty-one patients are eligible for this analysis. As expected, we observed a negative correlation between increases in proliferation and decreases in 45S CN (Figure S8, Spearman’s Rho = -0.35, P = 0.053; Fig S7B). Similarly, the data show a positive correlation between increases in proliferation and increases in the 5S / 45S ratio (Figure 7C, Rho = 0.43, P = 0.017; Fig S7B). The higher the increase in proliferation observed in a cancer lineage relative to the normal tissue within an individual the greater the loss in 45S rDNA and the higher the change in 5S/45S ratio. We also observed qualitatively similar pattern for LUSC and HNSC using both PRI sets (Table S5).

**Figure 7.**
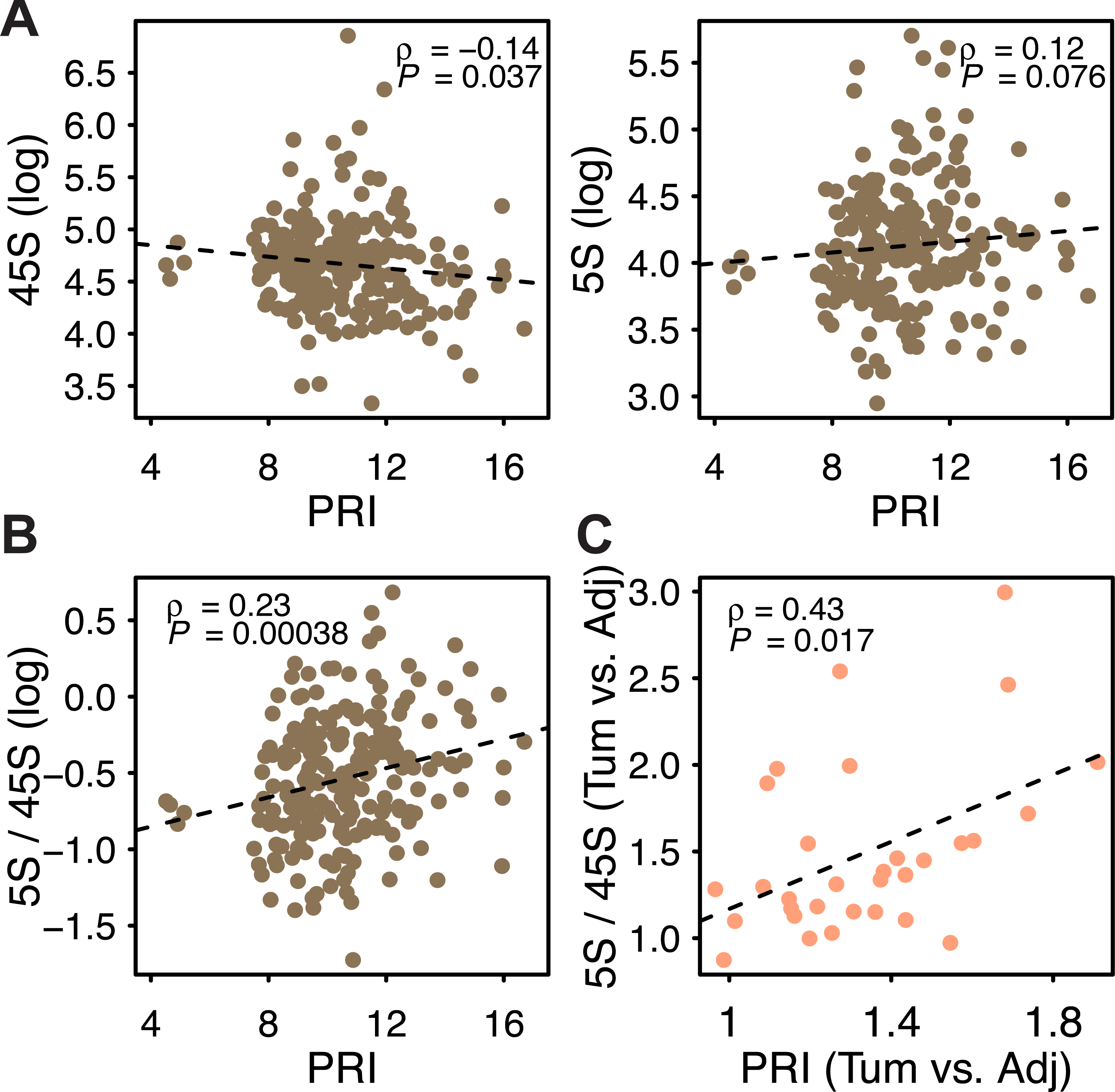
(A-B) Correlations between PRI (YW gene set) and 45S CN, 5S CN and 5S/45S ratio in LUAD tumor samples. (C) Changes in tumor PRI relative to adjacent tissue are positively correlated with changes in the 5S/45S ratio between tumor and adjacent tissue. The 31 LUAD patients with paired adjacent-tumor data were used in C.

### Discussion

Here we addressed variation in rDNA copy number across hundreds of individuals, tissue types, and within cancer lineages. The data revealed concerted copy number variation between the 5S and 45S rDNA arrays in some but not all tissue types. Furthermore, the data revealed coupled 5S rDNA amplification and 45S rDNA loss in cancer genomes relative to paired adjacent normal tissue, with rDNA changes that are associated with somatic mutations. For instance, coupled 5S gain / 45S loss are particularly salient in lineages with P53 inactivation from stomach and lung adenocarcinomas, but can also be observed in head and neck squamous cell carcinoma. Finally, we used global gene expression data to estimate tumor proliferation rates and show that they can partially explain coupled 5S rDNA gain and 45S rDNA loss.

It is important to highlight that the coupled events of 5S amplification and 45S loss that we observed in cancer linages is, on average, of small magnitude relative to the breadth of naturally occurring variation in 5S and 45S copy number that is observed across individuals in natural populations. For instance, ribosomal DNA copy number displays over 10-fold variation in LCLs from two human populations [3, 30]. Here we also observed a large breadth of variation in rDNA CN with 2-10 fold copy number changes among individuals for most tissues profiled. However, our comparisons between cancer and paired normal tissue indicate that more that 90% of tumors had less than 2-fold increases or decreases for either 5S or 45S rDNA. Similarly, most 1q42 amplification corresponded to duplications of the segment with less than 15% of the cases corresponding to higher increases (ploidy >= 2.5). How can rDNA copy number changes in cancer lineages be reconciled with large naturally occurring inter-individual variation in absolute rDNA copy number? One clue comes from the observation that somatic changes in the genetic context in cancers as well as changes in proliferation rates and ribosomal biogenesis influence rDNA CN. For instance, our analyses indicated that mutation in P53 is among the strongest determinants of coupled 5S gain and 45S loss in cancer lineages. Hence, the relative gain or loss of rDNA units is partially determined by somatic changes occurring within individuals. However, these somatic alterations are relatively minor when compared to genetic differences between individuals. The data raise the possibility that rDNA CN has a polygenic basis with inter-individual variation in rDNA array copy number strongly determined by genetic background, with increases and decreases within cancers reflecting rDNA adaptations to accelerate proliferation rates in tumor cells.

All aspects of rapid cell proliferation are dependent on efficient ribosomal biogenesis to sustain protein synthesis. Ribosomal RNA molecules are necessary structural components of eukaryotic ribosomes and high transcription rates from several rDNA units in the multicopy rDNA array are necessary to provide sufficient rRNAs for ribosome biogenesis. Indeed, up-regulation of ribosomal genes in cancer has been documented and is presumed to reflect the increased translational demand in rapidly proliferating cells [40, 52, 53]. Hence, the amplification of the 5S rDNA array in cancer is presumably selected by an increased demand for 5S rRNA molecules. It is also conceivable that lowered 45S rDNA CN could be adaptive and positively selected in cancer. This is because excess of rDNA copies has been suggested to promote global genome stability [35], and epigenetic regulation of the 45S rDNA could compensate for the loss of 45S rDNA alleles. If this is the case, the shorter 45S rDNA array in cancer could contribute to increased tumor evolvability through the promotion of genome instability. In this regard, drugs that target the rDNA might further fuel the adaptive ability of cancerous lineages. On the other hand, 45S rDNA loss might be a byproduct of replication stress emerging from both the rapid proliferation rates of cancer cells and the challenge of maintaining fast replication rates and high transcription rates in the 45S rDNA. Replication stress might be particularly salient in cancers lineages undergoing rapid cell cycle at the limit of their replication stress capacity. Indeed, transcription-replication conflict is common in eukaryotic genomes [54]. If cancer cells are indeed at the lower limit of frailty to balance rDNA replication, rDNA repair capacity, and rRNA transcription, drugs that target rDNA array copy number could be particularly effective.

Collectively, our observations about natural rDNA variation among genotypes and in cancer lineages suggest that gains or losses of rDNA units in cancer relative to normal adjacent tissue are more relevant in tumors than absolute rDNA copy number. The gain of 5S copies is expected to enhance ribosomal and nucleolar function in cancers whereas 45S copy number loss is a byproduct of replication-transcription conflict in rapidly proliferating cells. Interestingly, 45S rDNA loci are epigenetically regulated [55-57], with only a fraction of the alleles in an array being expressed at any time. Uncoupling 5S and 45S rDNA CN in tumors raise the possibility that cancers might be particularly reliant on epigenetic mechanisms to achieve higher 45S rRNA expression in the face of 45S rDNA loss. In this regard, drugs targeting epigenetic states of the rDNA could be especially effective. Our observations point to recurrent alteration in cancer cells, raise the prospects of using 5S and 45S ribosomal DNA states as indicators of cancer status and targets in new strategies for cancer therapy.

## Materials and Methods

### Obtaining the expression data

The RNAseq reads count of genes for different cancer types were downloaded from the Genomic Data Commons (GDC) data portal (https://gdc-portal.nci.nih.gov/), and normalized separately for each cancer type by implementing the ‘TMM’ method from the edgeR library in R[58]. RPKMs were calculated following standard procedures. We only included genes represented by at least 10 reads in more than half of all individuals.

### Selecting reference rDNA and background sequences

We used the 43kb consensus sequence of the 45S rDNA (GenBank accession: U13369.1). This sequence was modified to ~16kb (combining nucleotides from 41021-42999 with 1-14000), so that the transcription start site, the three mature components (18S, 5.8S and 28S), as well as the 3’ external transcribed spacer regions were all included. The 5S rDNA sequences and flanking regions was identified in an earlier study [3] and directly used here. From the genome annotation of Ensembl GRCh37.82 [59], we extracted exons and introns on chromosome 1, 13, 14, 15, 21 and 22 (where the rDNA arrays are located). If exons from multiple isoforms overlapped the largest isoform was selected; on the other hand, if an intron overlaps with an exon from a different isoform, then the intron is removed. We then performed similar filters as earlier studies [3, 30] to obtain groups of single copy exons and introns according to the following criteria. First, exons and introns having BLAST hit (e <10-6) [60] with any region (except for itself) of any gene, or any annotated human repetitive sequences from Repbase21.10 [61], were removed. Next, exons and introns with length smaller than 300 bps or larger than 10 kbs were removed. Finally, the first and last 50 nts of each region were not included. As a result, 2,453 exons and 3,091 introns were identified on chromosome 1, and 2,825 exons and 3,237 introns were identified on chromosome 13, 14, 15, 21, 22.

### Obtaining rDNA array reads

The mapped genome sequencing data (BAM files) of cancer patients generated by the TCGA project were downloaded from the Legacy Archive website of the GDC data portal (https://gdcportal.nci.nih.gov/legacy-archive/search/f). We used two approaches to obtain reads mapped onto rDNA sequences. In the first approach, BAM files were converted back to raw FASTQ reads using bamUtil v1.0.13 (https://github.com/statgen/bamUtil), and then mapped *de novo* onto the rDNA consensus sequences with BWA v0.7.9a [62]. These *de novo* mapping and rDNA estimates are similar to our earlier implementations [3, 30]. This approach could be quite computation consuming for samples with high sequencing depth. Alternatively, we noticed that (i) nearly all of the rDNA reads are mapped onto the GL000220.1 scaffold (for 45S) as well as 1q42 (for 5S) [3, 30], and that (ii) most of the samples had been mapped onto hg19 chromosomes plus the unintegrated scaffolds. Hence, we then used Samtools v1.3.1[63] to directly slice BAMs files and extract reads that had been aligned onto scaffold GL000220.1, and the 1q42 region (chr1:226743523-231781906. That is, up to 2MBs flanking both sides of the annotated 5S array, or chr1: 228743523- 228781906, were selected). After slicing, we aligned the subsets of reads onto the rDNA reference sequences. Note that multiple pseudo rRNA genes scatter across the genome. To account for the potential influence of these pseudo rRNAs for the first approach, we further mapped the rDNA reads onto human genome to calculate a correction ratio for each component, as suggested by earlier studies [3, 30]. However, it is not necessary to control for pseudo rRNAs in the slicing approach. We compared the results calculated from both approaches: only marginal differences were detected (Figure S2C). Hence, we adopted the slicing method throughout the study, because of its higher computational efficiency and comparable accuracy with *de novo* mapping.

### Estimation of background reads depth (BRD) for single copy exons and introns

From the downloaded BAM files, we calculated per base reads depth for the above groups of single copy exons and introns using the ‘depth’ command of Samtools. Note that cancer genomes suffer from large-scale structural variation, with frequent gain or loss of whole genes, partial segments as well as large-scale chromosomal duplications/losses. Therefore, we cannot assume that the selected background regions will necessarily maintain diploidy in cancer lineages. To account for the potential aneuploidy, we downloaded the gene-level copy number estimations produced as of January 28 2016 from the Broad FireBrowse portal. In this pipeline, genomic regions that undergo focal or arm-level amplification or deletion in tumors were identified by using the software GISTIC2 [43] from the human SNP array 6.0 datasets. One of the output files, ‘all_data_by_genes.txt’, contains gene-level copy number values (denoted as “C”) for each tumor sample. Here C = 0 means normal diploid, and 2 + C indicates the actual ploidy of the gene. We noticed a ranging ploidy for genes in tumor, supporting a need for correction (Figure S1). Ideally the ploidy of a gene corresponds with its sequencing depth. Encouragingly, we did observe strong positive correlations in tumors (Figure S2A). Therefore, we used a ratio, R = (2 + C)/2 to represent the fold change of gene copy in tumor relative to normal. We corrected the per-base coverage depth of selected exons and introns by the corresponding gene’s R-value in each tumor sample. The mean value of the adjusted depths was then used as BRD in tumors. On the other hand, BRD of adjacent or blood was computed as before without correction for ploidy [3, 30]. The correction process leads to improved rDNA CN estimates in tumors.

### Calculation of rDNA copy number

After obtaining reads mapped onto the rDNA references, we calculated the per-base depth for each component. The depths of 5.8S and 5S were averaged across all sites, a robust approach in view of the nearly identical depth estimates along these components. On the other hand, the two larger components, especially 28S, displayed considerable variability in depths across sites, which reflect underlying sequence polymorphism. To account for such heterogeneity, we considered two approaches. In the first approach, the mean depth is calculated as the average of the selected region of 18S (901-1871) or whole 28S sequence respectively. Alternatively, we examined a 150bp sliding-window in each component. For each 1 bp step we calculated the coefficient of variation (CV) of average depth across samples for the 150 nucleotides. The window with the lowest mean CV across LUAD adjacent samples were selected (18S:1145-1294, 28S:1522-1671). Moreover, we found that LUAD tumors have similar CV along 18S and 28S as adjacent, and that the selected windows also tend to have almost the least CV in LUAD tumors (Figure 1A), supporting the robustness of the selection. The same windows were applied for all other cancer types. Finally, to further obtain rDNA copy numbers, we divided the mean depth of 5S by the mean BRD of selected exons and introns from chromosome 1, whereas mean depth of 45S components were divided by BRD calculated from exons in chromosome 13, 14, 15, 21, 22, with corrections for aneuploidy in both cases. We also estimated CN for the overall 45S array by using the averaged depth across 18S, 5.8S and 28S components, with the slid windows applied for 18S and 28S.

### Identifying the plate ID effect and intra-plate variation

Plate ID (corresponding to sequencing center) of TCGA exome/genome sequencing data has been shown to affect CN estimates in previous studies [44, 64]. Hence, we carefully inspected our rDNA CN estimates to identify such effects. Specifically, since a number of samples were separately processed in different batches, we selected pairs of batches that shared at least 6 samples, and used paired Wilcoxon rank sum tests to identify potential differences between pairs from different batches. Ideally, shared samples should have similar CN estimates in each batch. However, as observed before, we did also observe significant differences for some comparisons (Figure S2B, Table S1). To account for such influence, we adopted the approach from [44]. Briefly, for each cancer type, a linear regression model was applied with tissue type and batch as independent variables, and logarithmic rDNA CN as the dependent variable. All 4 components, as well as the 45S array estimated above were corrected separately for each cancer type.

### Associating rDNA CN with somatic copy number alterations and non-silent mutations

To associate genome wide SCNAs with rDNA CN, we used significant local level amplification and deletion events identified in the FireBrowse portal. The extent of the SCNA is measured as “ploidy - 2” for amplification, and “2 - ploidy” for deletion. Tumors with a local ploidy change value larger than 0.1 (for amplifications) or smaller than -0.1 (for deletions) were identified as containing the SCNA. For each SCNA, we used a linear regression model to test the association between the SCNA event and the change in rDNA CN. Similarly, to associate somatic mutations with rDNA CN, we used “.maf” input file of the mutation analysis pipelines in the FireBrowse portal. We only included genes with non-silent somatic mutations in at least 10 patients. We used a Wilcoxon rank sum test to compare rDNA CN with in tumor lineages with and without the focal mutation.

## Calculating proliferation indexes (PRI)

We used two sets of independently identified genes to calculate PRI. The first set with 793 genes (or the ‘YW’ set) was obtained from Yedael Y. Waldman et al. [50], who calculated PRI from genes significantly positively correlated (Spearman’s Rho > 0, P < 0.01) with cell proliferation across 60 cancer cell lines. The second set of 350 genes (or the ‘RS’ set) was compiled by Rickard Sandberg et al. [51], who proposed the use of PRIs based on cell proliferation estimates obtained in non-cancerous cell lineages. Here we used the median expression of selected subset of genes to represent PRI. Interestingly, only 74 genes are shared between the two sets, yet both sets yielded highly correlated PRIs (Figure S4). To answer which cancers have significantly different PRI relative to adjacent tissue, the median FC of PRI genes between adjacent and tumor was calculated across patients. A minimum of 5 patients with paired adjacent and tumor RNAseq data for each cancer type were required.

## Availability of data and materials

The RNAseq processed data were downloaded from the GDC data portal (https://gdc-portal.nci.nih.gov/). The mapped genome sequencing data were downloaded from the Legacy Archive website of the GDC data portal (https://gdc-portal.nci.nih.gov/legacy-archive/search/f). The copy number variation and somatic mutation data were downloaded the FireBrowse portal (http://firebrowse.org/). All the data of cancer patients were generated or processed by the TCGA project (https://cancergenome.nih.gov/) and (http://www.ensembl.org/index.html). The two sets of genes used in the computation of the proliferation indexes were identified previously [50, 51].

## Acknowledgements

We gratefully acknowledge the TCGA consortium for the cancer data they generated. The computations in this paper were run on the Odyssey cluster supported by the Research Computing Group in the FAS Division of Science at Harvard University.

**The authors declare no conflicts of interest.**

## Legends to Supplementary Figures

**Figure S1.**
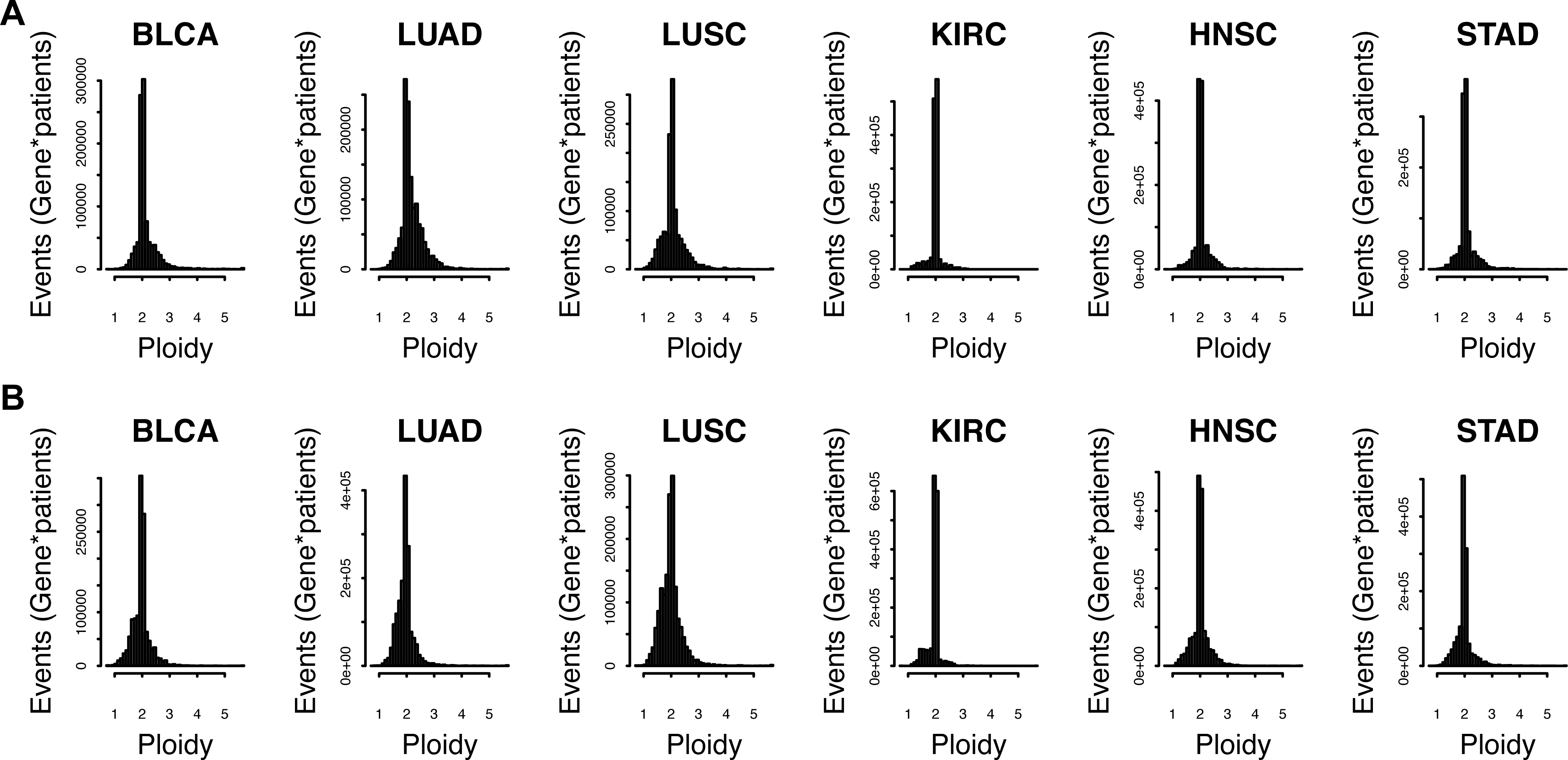
The distribution of ploidy for genes on (A) chromosome 1 and (B) five others (13, 14, 15, 21, 22) across cancers. Each gene from each tumor sample was regarded as an event.

**Figure S2.**
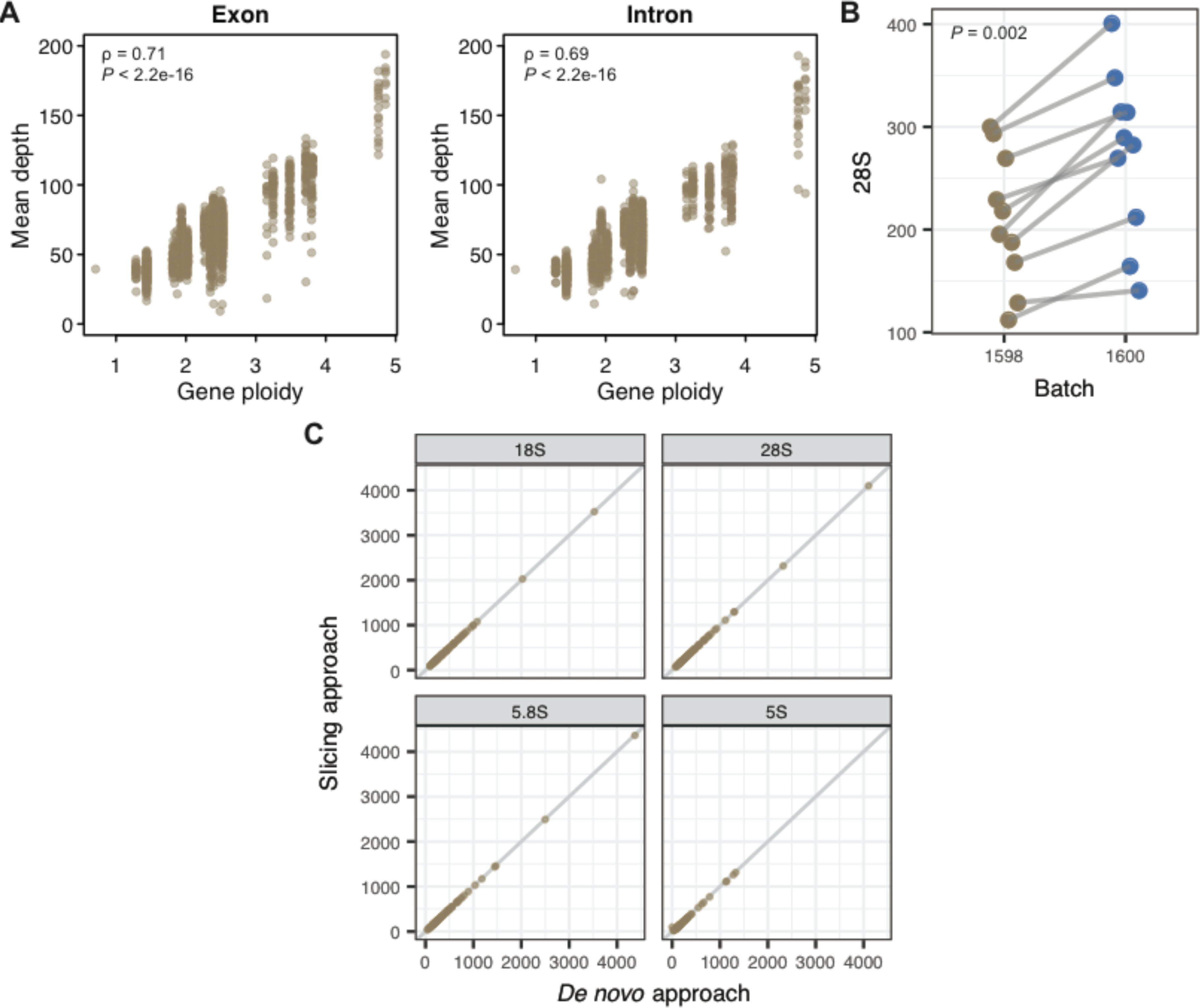
Estimating rDNA copy number in cancers. (A) Sequencing depth in exons (Spearman’s Rho = 0.71, P < 2.2e-16) and introns (Rho = 0.69, P < 2.2e-16) are strongly correlated with a gene’s ploidy in tumor. A LUAD tumor sample (TCGA-91-6847-01A-11D-1945-08) was randomly selected for this display. (B) Identical samples processed from batch 1600 had higher 28S copies than those of batch 1598 (Paired Wilcoxon rank sum test, P = 0.002). (C) Copy number (CN) estimates for all 4 components using two approaches (*de novo* mapping of raw reads or by “slicing” from pre-processed BAM files) are nearly identical across 100 randomly selected LUAD samples.

**Figure S3.**
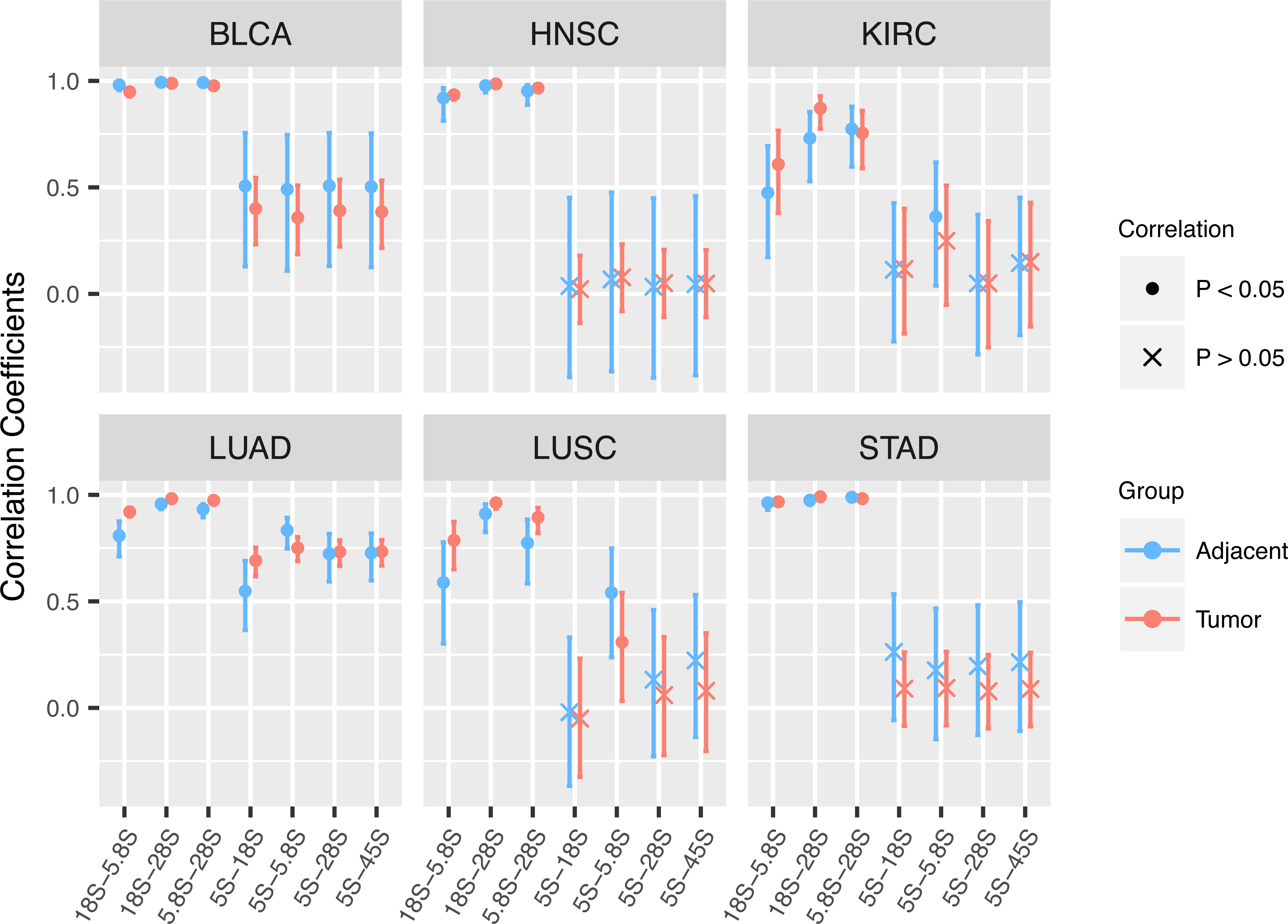
Pairwise correlation coefficients between rDNA components.

**Figure S4.**
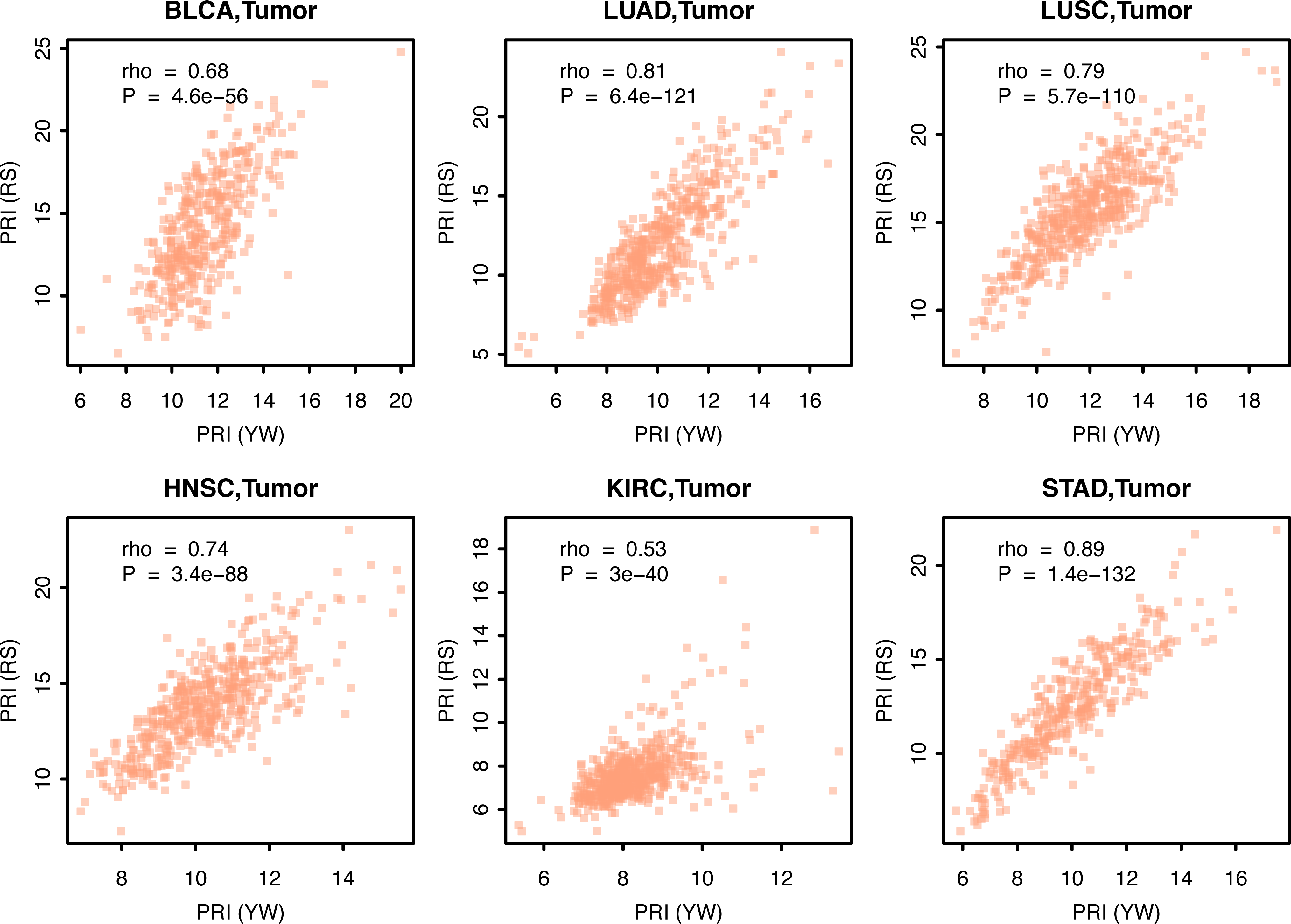
(A) PRIs calculated from the two independently defined gene sets are strongly correlated.

**Figure S5.**
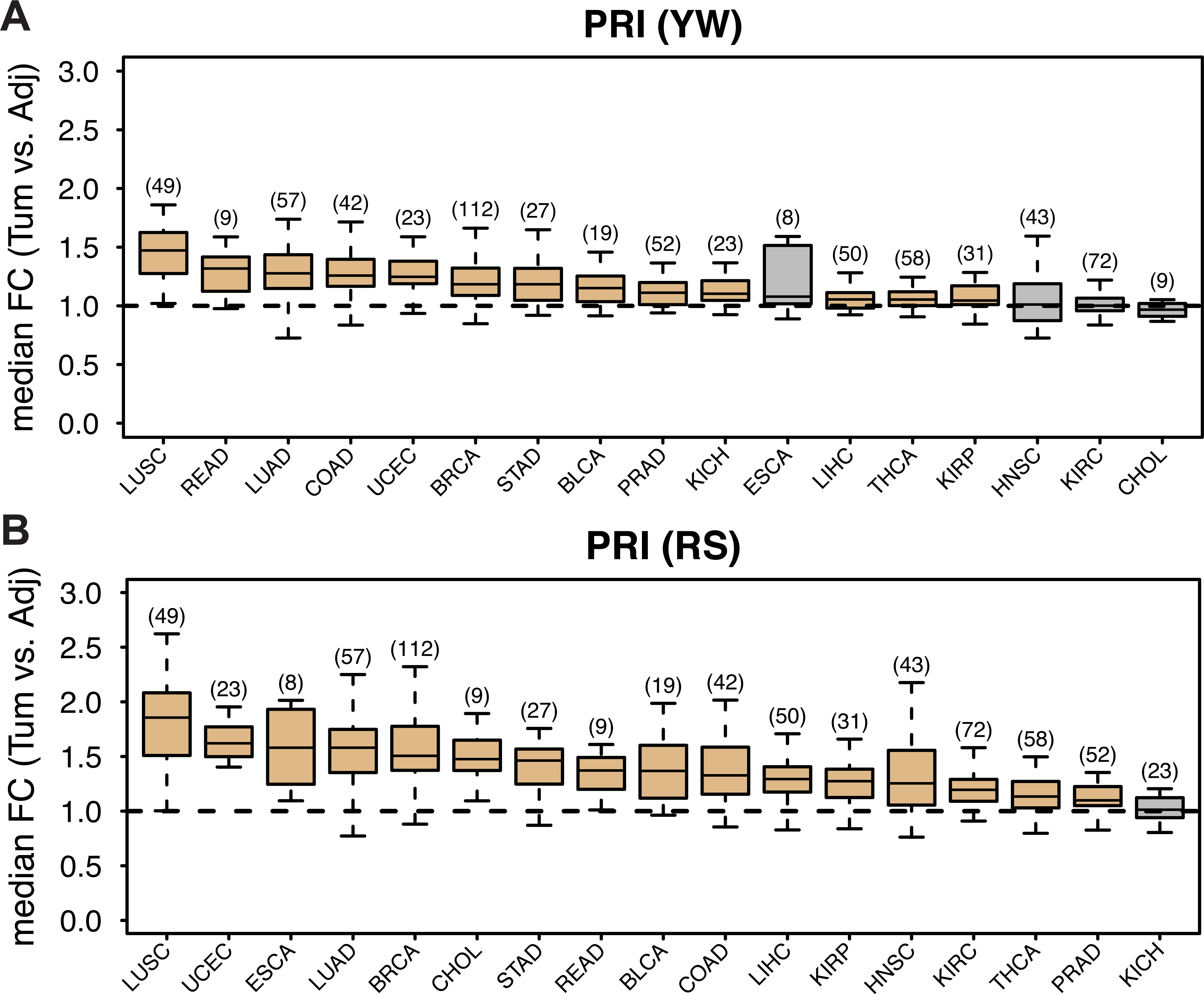
Most cancer types display increased proliferation index (PRI). PRI were calculated from (A) YW set and (B) RS set. Yellow and grey indicate significant upregulation and not significant in tumors compared with paired adjacent controls (Wilcoxon rank sum test P < 0.01), respectively. Sample sizes are indicated in brackets. UCEC, uterine corpus endometrial carcinoma; ESCA, esophageal carcinoma; CHOL, cholangiocarcinoma; READ, rectum adenocarcinoma; COAD, colon adenocarcinoma; LIHC, liver hepatocellular carcinoma; KIRP, kidney renal papillary cell carcinoma; THCA, thyroid carcinoma; PRAD, prostate adenocarcinoma; KICH, kidney chromophobe. Other abbreviations are as in Table 1.

**Figure S6.**
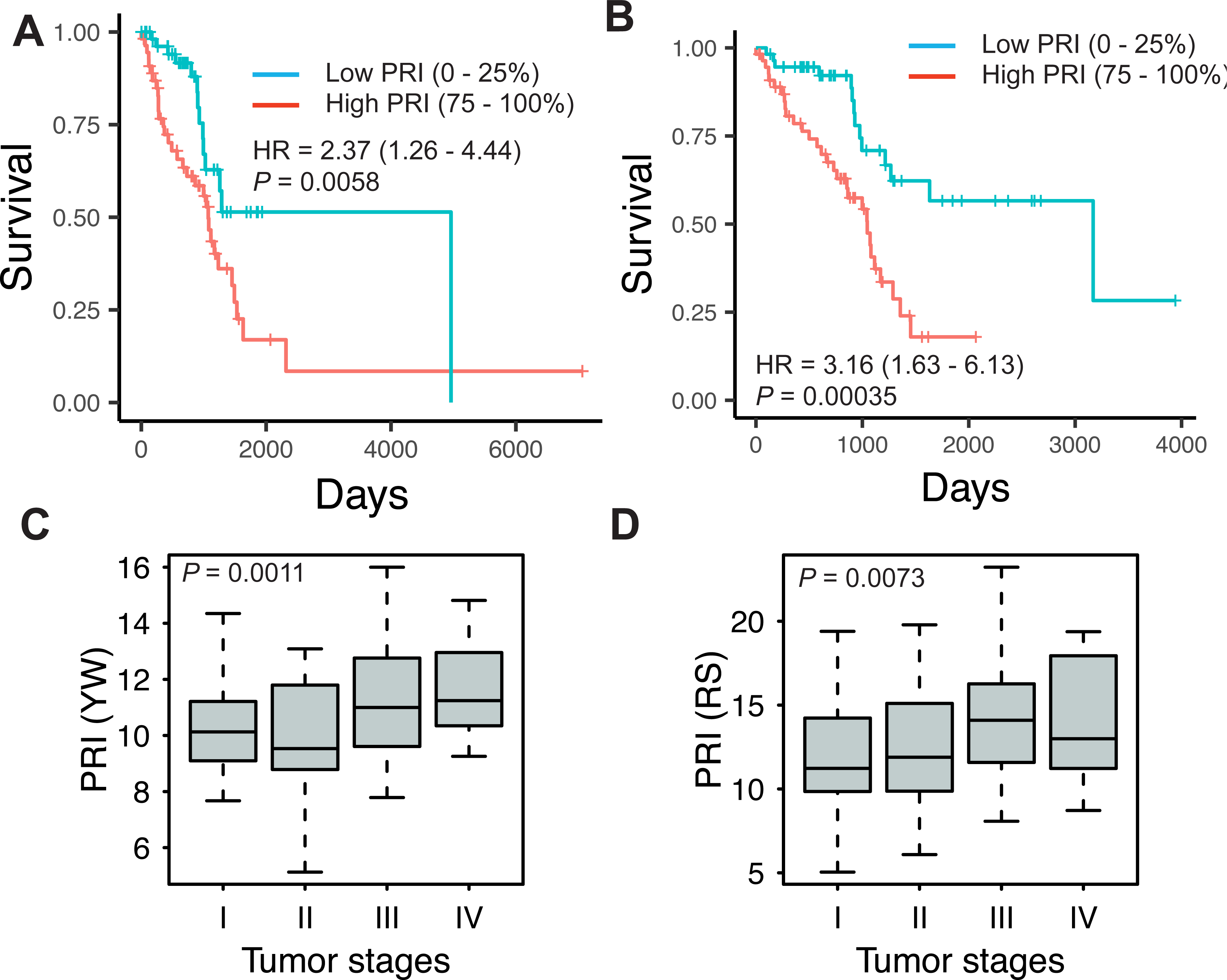
Example using LUAD data showing that patients with higher PRI tend to have (A, B) worse survival (comparing the last with the first 25% patients, logrank test, P < 0.006, Hazards ratio > 2.35), as well as (C, D) more severe tumor stage (ANOVA, P < 0.0075). The YW set genes are used for A and C; while the RS set genes are used for B and D.

**Figure S7.**
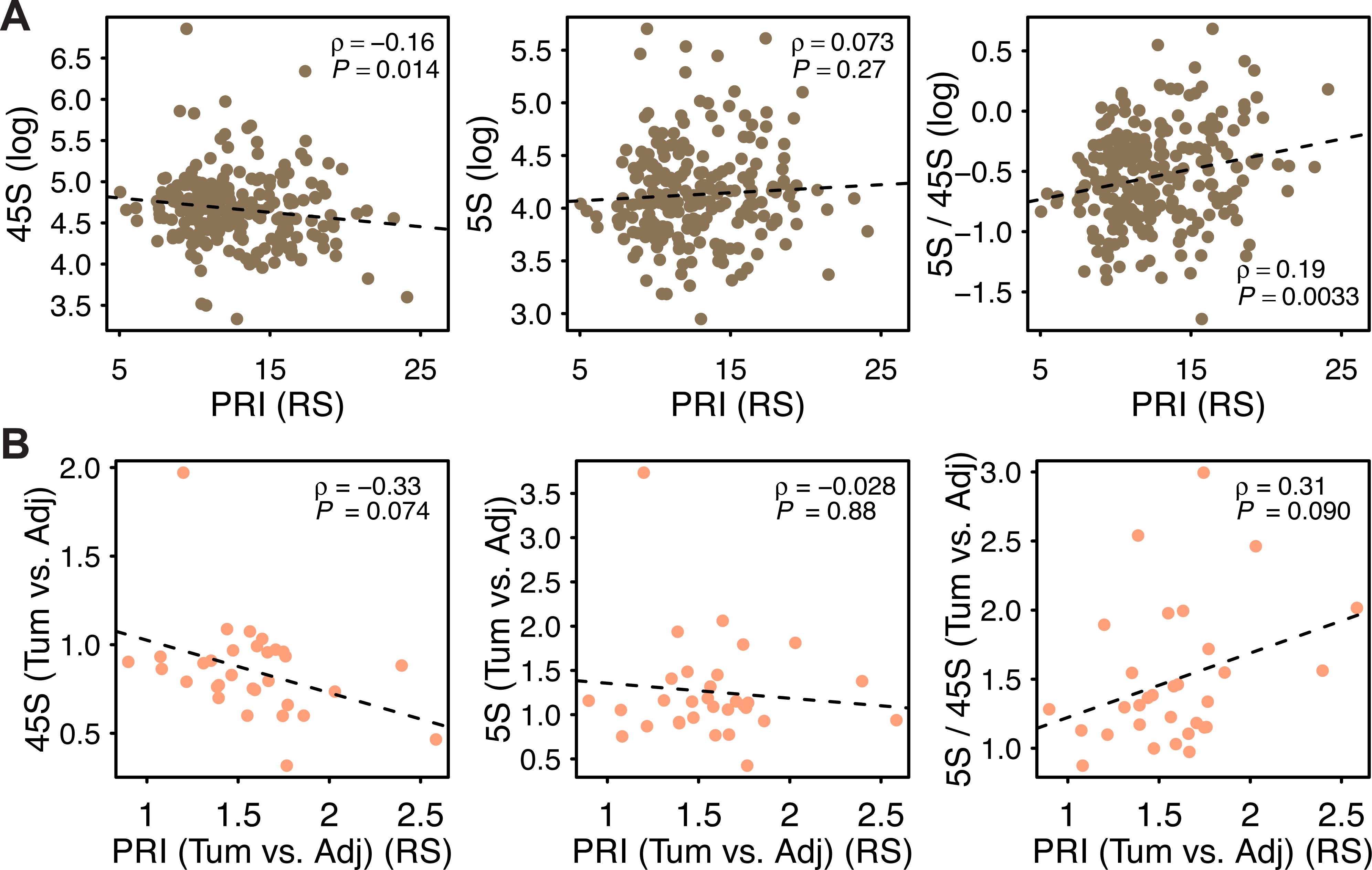
(A) PRI is significant negatively correlated with 45S and positively with the 5S / 45S ratio whereas it is not significant with 5S in tumors. (B) Consistent results were observed when associating tumor vs. adjacent fold change of PRI with that of 5S, 45S or their ratio for 31 patients. RS set genes and LUAD samples are used.

**Figure S8.**
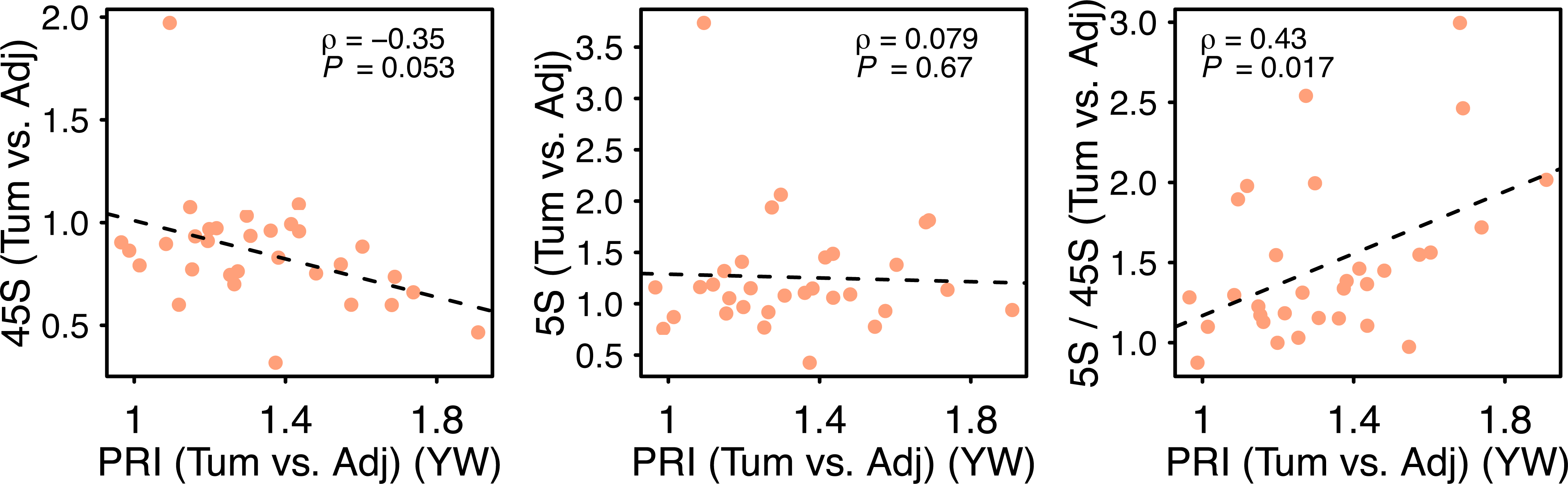
The correlation between relative fold change of proliferation (YW set) in tumor relative to its adjacent, and the fold change of 5S, 45S or their ratio for same patients. LUAD samples are used.

## Legends to Supplementary Tables

**Table S1.** Sequencing batch (plate ID) effects on rDNA CN estimation. Samples sequenced on two separate plates (plate 1 and plate 2) were identified. These were selected and their CN compared with a Wilcox rank sum test.

**Table S2.** Association between rDNA CN with somatic copy number alternations (SCNAs)

**Table S3.** Association between rDNA CN with somatic mutations. Each cancer type was analyzed separately.

**Table S4.** Association between rDNA CN with somatic mutations. All available cancers were pooled.

**Table S5.** Association between rDNA CN and proliferation index (PRI) in tumors

